# Developmental and conditional regulation of DAF-2/INSR ubiquitination in *Caenorhabditis elegans*

**DOI:** 10.1101/2024.05.24.595723

**Authors:** Ivan B. Falsztyn, Seth M. Taylor, L. Ryan Baugh

## Abstract

Insulin/IGF signaling (IIS) regulates developmental and metabolic plasticity. Conditional regulation of insulin-like peptide expression and secretion promotes different phenotypes in different environments. However, IIS can also be regulated by other, less-understood mechanisms. For example, stability of the only known insulin/IGF receptor in *C. elegans*, DAF-2/INSR, is regulated by CHIP-dependent ubiquitination. Disruption of *chn-1/CHIP* reduces longevity in *C. elegans* by increasing DAF-2/INSR abundance and IIS activity in adults. Likewise, mutation of a ubiquitination site causes *daf-2(gk390525)* to display gain-of-function phenotypes in adults. However, we show that this allele displays loss-of-function phenotypes in larvae, and that its effect on IIS activity transitions from negative to positive during development. In contrast, the allele acts like a gain-of-function in larvae cultured at high temperature, inhibiting temperature-dependent dauer formation. Disruption of *chn-1/CHIP* causes an increase in IIS activity in starved L1 larvae, unlike *daf-2(gk390525)*. CHN-1/CHIP ubiquitinates DAF-2/INSR at multiple sites. These results suggest that the sites that are functionally relevant to negative regulation of IIS vary in larvae and adults, at different temperatures, and in nutrient-dependent fashion, revealing additional layers of IIS regulation.

**ARTICLE SUMMARY:** Insulin-like signaling plays a critical role in helping animals adapt to different environmental conditions. Differences in abundance of insulin molecules drive differences in insulin signaling, affecting growth, metabolism, and resistance to stressful conditions. Previous work in the roundworm *C. elegans* showed that targeted degradation of the insulin receptor also regulates insulin signaling. We show here that this process is affected by developmental stage, nutrient availability, and temperature, revealing additional ways that insulin-like signaling is regulated in this valuable animal model.

## INTRODUCTION

IIS regulates growth, development, metabolism, stress resistance, and aging in metazoans. In the nematode *C. elegans*, the sole known insulin/IGF receptor is encoded by *daf-2/INSR* (Murphy and Hu 2013). DAF-2/INSR signals through a conserved phosphoinositide 3-kinase (PI3K) pathway including AGE-1/PI3K, PDK-1, AKT-1, and AKT-2 to govern nuclear localization and activity of the transcription factor DAF-16/FOXO.

When worms hatch in the absence of food, they remain in a developmentally arrested state in the first larval stage known as L1 arrest (or L1 diapause) (Baugh 2013). Extended starvation during L1 arrest causes developmental abnormalities of the gonad, including germline tumors (Jordan *et al*. 2019). Worms also arrest development as dauer larvae in the third larval stage in response to adverse environmental conditions including high population density, limited nutrient availability, and high temperature (Hu 2007). IIS regulates L1 arrest and dauer formation, with *daf-2/INSR* mutants displaying constitutive arrest phenotypes as L1 and dauer larvae (Gems *et al*. 1998; Baugh and Sternberg 2006) as well as increased starvation resistance, including increased survival of L1 arrest (Muñoz and Riddle 2003; Baugh and Sternberg 2006) and suppression of starvation-induced developmental abnormalities (Jordan *et al*. 2019). IIS also regulates adult physiology, and disruption of *daf-2/INSR* substantially increases lifespan and suppresses vitellogenesis (Cynthia Kenyon *et al*. 1993; Murphy *et al*. 2003; Depina *et al*. 2011).

Ubiquitin is a small peptide that is covalently attached to proteins. Ubiquitination often targets proteins for degradation, but it can regulate protein function in other ways as well (Kipreos 2005; Zheng and Shabek 2017). The quality-control E3 ubiquitin ligase CHIP mono-ubiquitinates worm, fly, and human INSR, with DAF-2/INSR being ubiquitinated at multiple sites (Tawo *et al*. 2017). DAF-2/INSR stability is regulated by the sole worm ortholog of CHIP, encoded by gene *chn-1*. Disruption of *chn-1/CHIP* increases DAF-2/INSR abundance in adults, increasing IIS activity and reducing lifespan (Tawo *et al*. 2017). *daf-2(gk390525)* is a point mutation resulting in a K1614E amino acid substitution, affecting one of the lysine residues targeted by CHN-1/CHIP-dependent ubiquitination (Tawo *et al*. 2017). This mutant has reduced lifespan (Tawo *et al*. 2017; Zhao *et al*. 2021) and increased vitellogenesis (Kern *et al*. 2021), consistent with increased IIS activity, causing it to be described as a gain-of-function allele. Although the effects of *chn-1/CHIP* and DAF-2/INSR ubiquitination have been documented in adults, it is unknown if or how ubiquitination affects IIS during larval development or in conditions where there are large differences in IIS, such as in fed vs. starved animals.

We characterized *daf-2(gk390525)* and *chn-1(by155)* phenotypes in L1 arrest and recovery, dauer formation, and in adults. We complemented phenotypic analysis with quantification of DAF-16/FOXO sub-cellular localization as a proxy for IIS activity. We confirm published results showing that this allele causes increased IIS activity in adults. We also show that it increases IIS activity during temperature-dependent dauer formation. However, we demonstrate that it reduces IIS activity during larval development and L1 arrest, behaving like a loss-of-function allele. In contrast to *daf-2(gk390525)*, disruption of *chn-1/CHIP* increases IIS during L1 arrest, suggesting regulation of DAF-2/INSR through ubiquitination of one or more amino acid residues other than K1614. These results demonstrate that *chn-1/CHIP* and ubiquitin-dependent regulation of DAF-2/INSR varies during development and in different conditions. They also highlight that *daf-2(gk390525)* has complex effects on IIS, warranting caution in interpretation of phenotypic analysis.

## RESULTS AND DISCUSSION

### *daf-2(gk390525)* behaves as a gain-of-function allele in adults

We assayed lifespan for *daf-2(gk390525)*, a reportedly gain-of-function allele (Tawo *et al*. 2017; Kern *et al*. 2021; Zhao *et al*. 2021), *daf-2(e1370)*, a class 2 loss-of-function allele (Gems *et al*. 1998), and *daf-18(ok480)*, a null allele of a negative regulator (*daf-18/PTEN*) of AGE-1/PI3K and thus IIS (Ogg and Ruvkun 1998). As expected, *daf-2(e1370)* had a significant increase in lifespan relative to the wild-type (N2) control, and *daf-18(ok480)* had a significant decrease (Fig. 1a). *daf-2(gk390525)* had a relatively small, albeit significant, decrease in lifespan. These results confirm the published gain-of-function behavior of *daf-2(gk390525)* with respect to lifespan (Tawo *et al*. 2017; Zhao *et al*. 2021).

**Figure 1.**
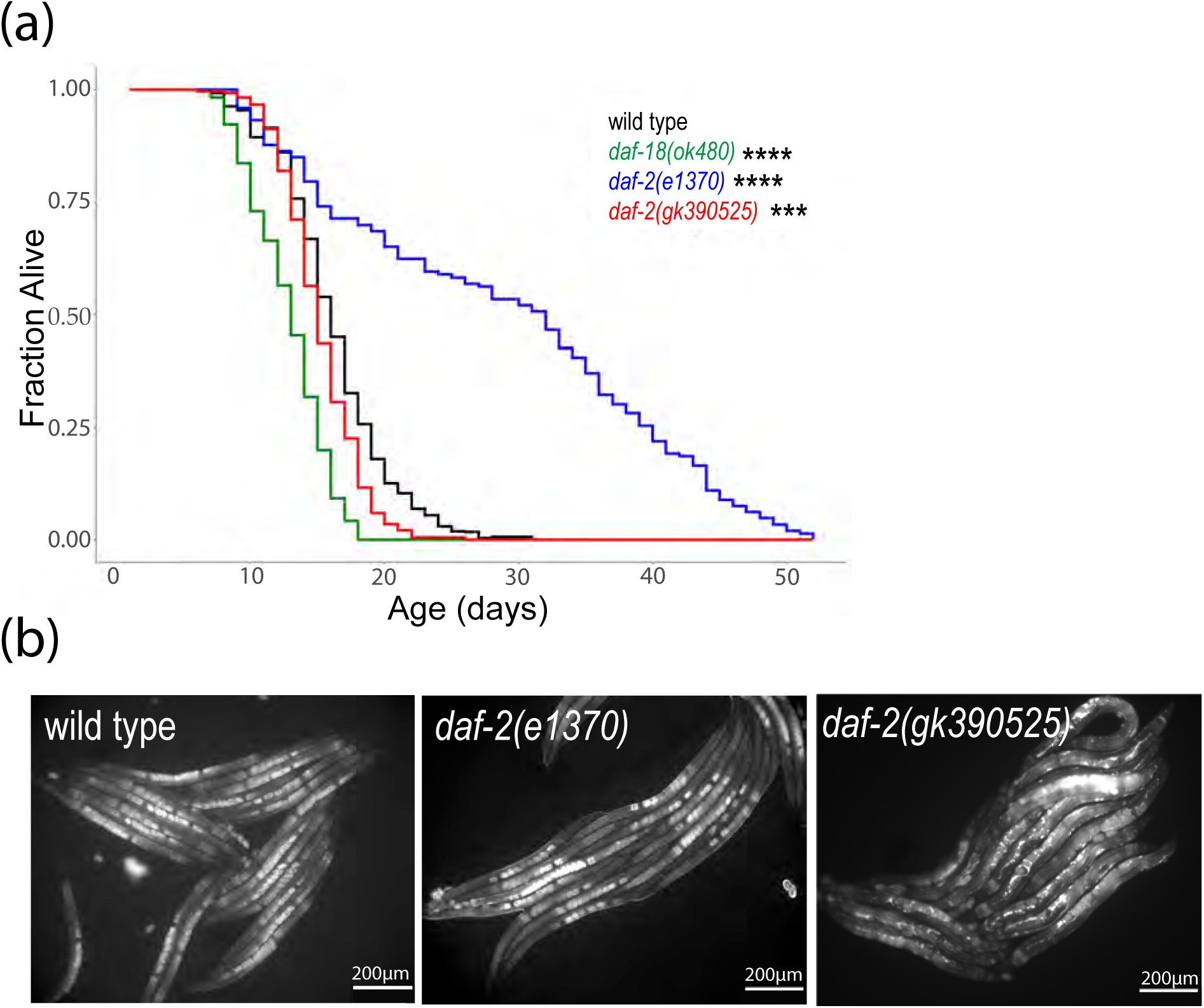
*daf-2(gk390525)* mutants demonstrate documented IIS gain-of-function phenotypes in adults. A) Adult lifespan was scored daily for three or five biological replicates, and pooled results are presented. See Table S1 for complete results. *** P < 0.001, **** P < 0.0001; log-rank test. B) Representative images of VIT-2::GFP expression were taken in the indicated genetic backgrounds at 200x total magnification. Well-fed animals were imaged during early adulthood, when a single row of embryos was visible in the uterus. To ensure matching stages, wild type and *daf-2(e1370)* were imaged after 84 hours recovery from L1 arrest, and *daf-2(gk390525)* was imaged after 72 hours.

DAF-16/FOXO antagonizes vitellogenesis (Murphy *et al*. 2003; Depina *et al*. 2011), and IIS promotes vitellogenesis and yolk venting (Kern *et al*. 2021). We imaged expression of a multi-copy VIT-2 reporter gene in *daf-2(e1370)* and *daf-2(gk390525)* gravid adults. VIT-2 and other vitellogenin proteins are synthesized in the intestine and secreted into the body cavity, and oocytes are provisioned through receptor-mediated endocytosis of vitellogenin lipoprotein particles (Perez and Lehner 2019). VIT-2::GFP expression appeared lower in the intestine and body cavity of *daf-2(e1370)* compared to wild type (Fig. 1b), consistent with reduced IIS and vitellogenesis. However, embryos *in utero* appeared brighter in *daf-2(e1370)*, consistent with increased vitellogenin provisioning with reduced IIS (Jordan *et al*. 2019). In contrast, *daf-2(gk390525)* appeared to have a higher total signal for VIT-2::GFP, with expression conspicuously increased in the body cavity. These results confirm the published gain-of-function behavior of *daf-2(gk390525)* with respect to vitellogenesis (Kern *et al*. 2021).

### *daf-2(gk390525)* behaves as a loss-of-function allele in fed and starved larvae

DAF-2/INSR and IIS affect a variety of larval phenotypes, but the effect of *daf-2(gk390525)* on them is unknown. We assayed multiple phenotypes related to L1 arrest and recovery in *daf-2(gk390525)* and *daf-2(e1370)* mutants. IIS regulates survival during L1 arrest. *daf-2(e1370)* survived L1 arrest significantly longer than wild-type (Fig. 2a), as expected (Muñoz and Riddle 2003; Baugh and Sternberg 2006; Hibshman *et al*. 2017). However, *daf-2(gk390525)* also displayed a relatively minor but significant starvation-resistant phenotype, suggesting loss of function during L1 arrest, despite displaying gain-of-function behavior in adults (Fig. 1).

**Figure 2.**
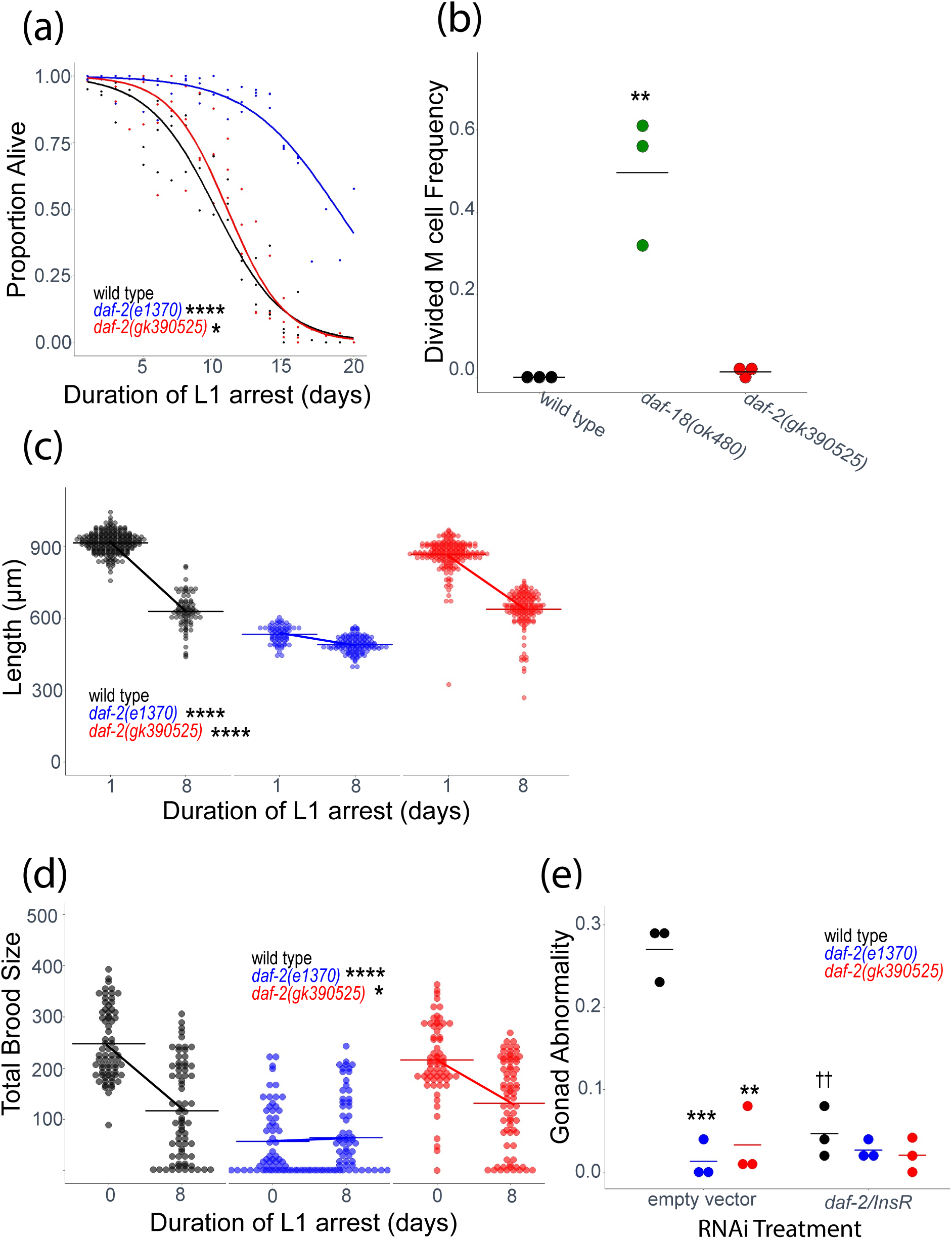
*daf-2(gk390525)* mutants display IIS loss-of-function larval phenotypes related to L1 starvation resistance. A) L1 starvation survival was scored daily for three biological replicates. Individual points represent an observation for a population of ∼100 animals (median = 97, range = 36-152) in a single biological replicate, and curves were fit with logistic regression. Two-tailed, unpaired variance t-tests were used to compare half-lives between wild type and each mutant. See Table S2 for complete data. B) M cell divisions were scored after 3 days of L1 arrest in the *daf-18(ok480)* mutant background, and after 8 days for all other genotypes using an *hlh-8* reporter gene as an M cell marker in three biological replicates. Horizontal lines represent the mean proportion of animals with at least one M cell division across replicates, and points represent the proportion per replicate (scoring ∼100 animals). A one-sided t-test was used to assess significance. See Table S2 for complete survival data. C) Worms were imaged and body length was measured after 48 hours of recovery from 1 (control – arrested L1s plated 24 hr after hypochlorite treatment) or 8 d L1 arrest in three biological replicates. Individual points represent an observation for a single animal. D) Total brood size was scored for ∼18 individual worms in each of three biological replicates (median = 18, range = 11-18). Worms recovered from 8 d L1 arrest are compared to unstarved controls (0 days L1 arrest – embryos plated directly with food). Individual points represent individual animals. E) The frequency of adults with gonad abnormalities was scored in previously starved animals (8 days L1 arrest) in three biological replicates. Individual points represent an observation in a population of ∼50 individuals in a single biological replicate. Horizontal lines represent the mean abnormality frequency across replicates within the same condition. Homogeneity of variance across conditions and replicates was tested using Bartlett’s test, and if variances were found to not be unequal they were pooled for statistical analysis. Two-tailed, unpaired t-tests with variance pooled were used to compare the frequency of gonad abnormalities between each mutant and wild type (indicated with asterisks) and between each RNAi treatment and empty vector control (indicated with crosses). C, D) A two-factor, linear, mixed-effects model was fit to the data with body length (C) or total brood size (D) as the response variable, interaction between starvation and genotype as the fixed effects, and replicate as the random effect. P-values for interaction terms are reported, assessing starvation-dependent effects of each genotype on phenotype. Horizontal lines represent the mean length per condition across replicates, and diagonal lines connecting the means between conditions within the same genotype represent the effect of starvation. A-E) * P < 0.05, ** P < 0.01, *** P < 0.001, **** P < 0.0001, †† P < 0.01.

In wild-type larvae hatched in the absence of food, the M cell does not divide, reflecting developmental arrest (Fig. 2a; Baugh & Sternberg, 2006). In contrast, there were substantial M cell divisions in *daf-18(ok480)* mutants, as expected with increased IIS (Chen *et al*. 2022). Over-expression of agonistic insulin-like peptides during L1 starvation drives M cell division (Chen and Baugh 2014), but there were very few M cell divisions in *daf-2(gk390525),* consistent with this allele not appreciably increasing IIS during L1 arrest (Fig. 2b).

Development is delayed following L1 arrest (Jobson *et al*. 2015), but reduction of IIS mitigates delay (Olmedo *et al*. 2020). We used image analysis to assay larval length after 48 hr recovery from L1 arrest. *daf-2(e1370)* larvae were smaller than wild-type in control conditions (1 d L1 arrest, which is valuable for synchronization), as expected, but there was very little additional effect of 8 d of L1 arrest, reflecting substantial starvation resistance (Fig. 2c). Length of *daf-2(gk390525)* larvae was only modestly affected in control conditions, and there was a substantial effect of 8 d of L1 arrest on size, but the effect of starvation was significantly dampened compared to wild type, again suggesting loss of function.

Brood size is reduced following extended L1 arrest (Jobson *et al*. 2015), but disruption of *daf-2/INSR* function mitigates decreased fecundity (Jordan *et al*. 2019). *daf-2(e1370)* brood size was substantially reduced in control conditions (no starvation), as expected, but brood size was not affected by 8 d L1 arrest, again reflecting substantial starvation resistance (Fig. 2d). *daf-2(gk390525)* brood size was only modestly affected in control conditions, but the effect of starvation was significantly dampened compared to wild type, further suggesting loss of function.

Extended L1 arrest leads to development of germline tumors and other developmental abnormalities in the adult gonad, and disruption of *daf-2/INSR* suppresses formation of these starvation-induced abnormalities (Jordan *et al*. 2019, 2023; Shaul *et al*. 2022). *daf-2(e1370)* and *daf-2* RNAi significantly suppressed formation of starvation-induced abnormalities (Fig. 2e), as expected. *daf-2(gk390525)* also significantly suppressed development of gonad abnormalities. In addition, *daf-2* RNAi of *daf-2(gk390525)* larvae during recovery from L1 arrest had no effect. These observations further support the conclusion that *daf-2(gk390525)* functions as a loss-of-function allele during L1 arrest and possibly recovery.

DAF-16/FOXO is the primary transcriptional effector of IIS and is antagonized by DAF-2/INSR and AGE-1/PI3K signaling (Murphy and Hu 2013). When IIS is active (e.g., in fed larvae), DAF-16/FOXO is phosphorylated and cytoplasmic, but when IIS is reduced (e.g., in starved larvae), DAF-16/FOXO translocates to the nucleus and regulates transcription (Henderson and Johnson 2001). We assayed DAF-16::GFP sub-cellular localization as a proxy for IIS activity to complement phenotypic analysis. DAF-16::GFP nuclear localization was significantly increased in starved L1 larvae compared to fed L1 larvae (Fig. 3a), as expected. In addition, DAF-16::GFP was almost entirely cytoplasmic and almost entirely nuclear in both conditions in *daf-18(ok480)* and *daf-2(e1370)* larvae, respectively, as expected for constitutively increased and decreased IIS. DAF-16::GFP displayed a modest but significant increase in nuclear localization in *daf-2(gk390525)* starved L1 larvae compared to fed (Fig. 3a), suggesting reduced environmental responsiveness compared to wild type. Moreover, DAF-16::GFP was significantly more nuclear in *daf-2(gk390525)* fed and starved L1 larvae compared to wild type in each condition, supporting the conclusion that it reduces DAF-2/INSR function in L1 larvae. Notably, these results extend the phenotypic analysis of L1 arrest and recovery (Fig. 2) to show that *daf-2(gk390525)* behaves as a loss-of-function allele in starved and fed L1 larvae.

**Figure 3.**
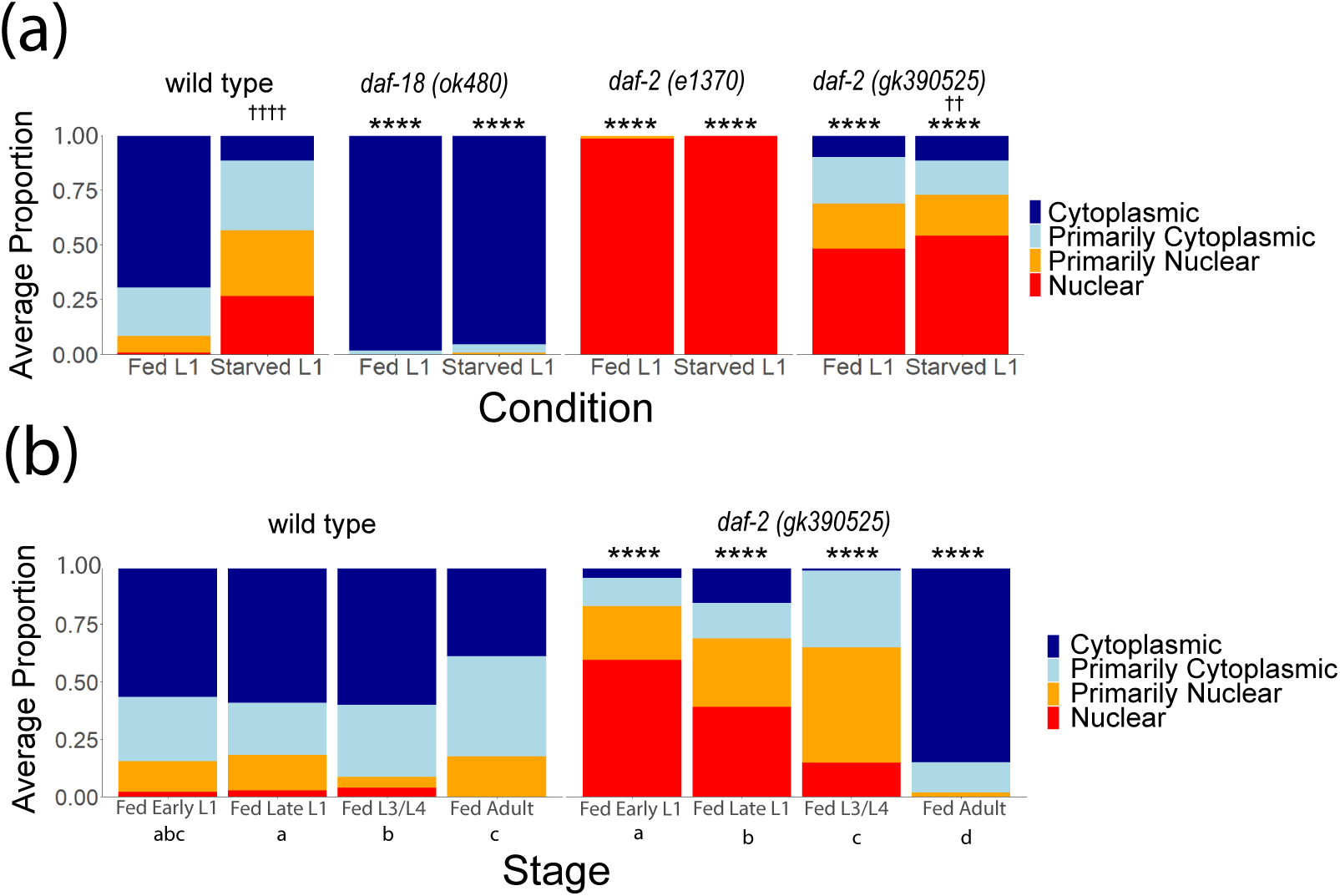
DAF-16 localization in *daf-2(gk390525)* reveals a developmental transition from loss-of-function behavior in larvae to gain-of-function behavior in adults. A) DAF-16::GFP sub-cellular localization was scored in fed (18 hr after hypochlorite treatment, or about 6 hr L1 development) and starved (24 hr after hypochlorite treatment, or about 12 hr L1 arrest) L1 larvae. Significant differences within genotype between conditions are indicated with crosses. B) DAF-16::GFP sub-cellular localization was scored throughout postembryonic development (18, 24, 48, and 72 hr after hypochlorite treatment, or about 6, 12, 36, and 60 hr postembryonic development, respectively). Statistically significant differences between stages within the same genotype are indicated with letters (a, b, and c), with members of groups sharing letters not being significantly different from each other (p = 0.05). A, B) Localization was scored as nuclear, primarily nuclear, primarily cytoplasmic, or cytoplasmic in intestinal cells in three biological replicates. Average frequency across replicates for each category is displayed. Cochran-Mantel-Haenszel chi-squared tests were used to compare the distribution of cellular localization between genotypes and conditions. Comparisons between each mutant and wild type within the same conditions are indicated with asterisks. ** P < 0.01, **** P < 0.0001. †† P < 0.01, †††† P < 0.0001

### Disruption of DAF-2/INSR ubiquitination at K1614 has different effects on IIS in larvae and adults

*daf-2(gk390525)* behaves as a gain-of-function allele in adults but a loss-of-function in L1 larvae, leading us to hypothesize that its functional impact on IIS transitions during development. To test this hypothesis, we scored DAF-16::GFP localization over the course of larval development. Fed wild-type animals displayed relatively minor fluctuations in localization throughout development, with DAF-16::GFP being mostly cytoplasmic (Fig. 3b), as expected. In contrast, DAF-16::GFP was primarily nuclear in *daf-2(gk390525)* fed early L1 larvae, as previously observed (Fig. 3a). In fed late L1 larvae, DAF-16::GFP shifted towards being more cytoplasmic compared to early L1 larvae (Fig. 3b). Likewise, a further shift towards cytoplasmic localization was observed in L3/L4 larvae, and again in adults. Critically, DAF-16::GFP was significantly more nuclear in mutant early L1, late L1, and L3/L4 larvae compared to wild-type, but it was significantly more cytoplasmic in adults. These results demonstrate that the effect of *daf-2(gk390525)* gradually shifts during development from reducing to increasing IIS, with the transition from behaving as a loss-of-function to gain-of-function allele occurring near the onset of adulthood.

### Disruption of DAF-2/INSR ubiquitination at K1614 has different effects on IIS at different temperatures

Unlike L1 arrest, dauer arrest occurs in the L3 stage in larvae that had at least some food (enough to develop into dauer larvae) but experienced adverse conditions such as high population density, limited nutrient availability, and/or high temperature (Hu 2007). Multiple signaling pathways regulate dauer formation, including IIS. In conditions of high IIS, larvae proceed through larval development into reproductive adults, but with low IIS they arrest as dauer larvae. Given the ecological significance of dauer development and the fact that dauer formation is one of the most profound consequences of reduced IIS, we wondered how *daf-2(gk390525)* affects dauer formation. The comfort range for *C. elegans* development is 15°C to 25°C, and wild-type larvae can develop as dauers at 27°C. We did not observe dauer formation in wild type, *daf-18(ok480)*, *daf-2(e1370)*, or *daf-2(gk390525)* at 20°C (Fig. 4a), as expected. However, at 25°C *daf-2(e1370)* displayed significant dauer formation, as expected, though *daf-2(gk390525)* did not. At 27°C, wild type displayed modest but reproducible dauer formation and *daf-2(e1370)* formed 100% dauers, as expected. However, *daf-18(ok480)* did not form dauers at 27°C, consistent with increased IIS, as expected. Likewise, dauer formation was significantly suppressed at 27°C in *daf-2(gk390525)* compared to wild type, suggesting increased IIS. These results suggest that *daf-2(gk390525)* behaves like a gain-of-function allele in larvae at 27°C, despite its loss-of-function behavior in larvae at 20°C.

**Figure 4.**
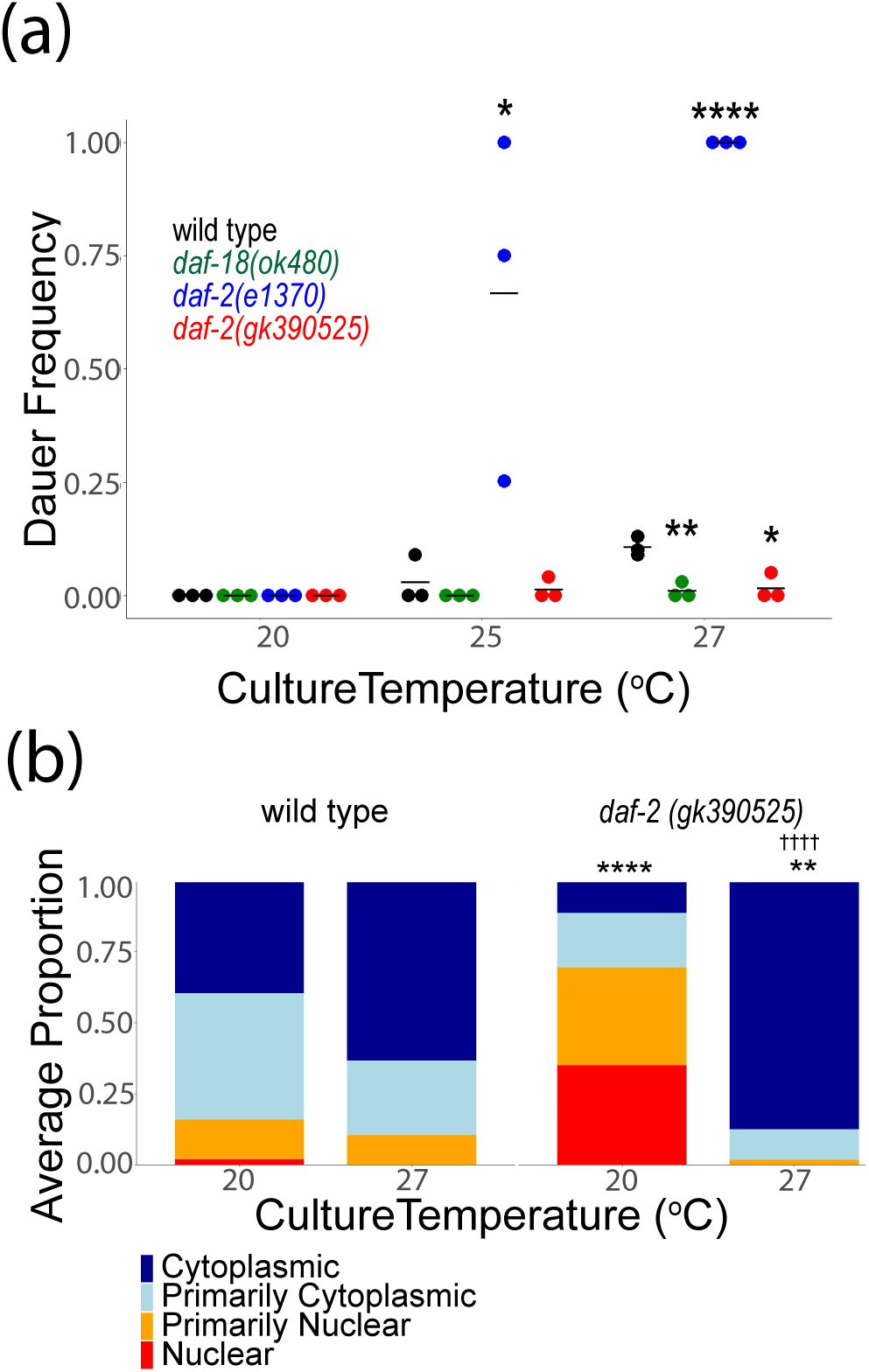
*daf-2(gk390525)* mutants display IIS gain-of-function phenotypes in larvae at elevated temperature. A) The frequency of dauer larvae was scored at two different temperatures in three biological replicates. Each individual point represents an observation for a population ∼100 individuals in a single biological replicate. Horizontal lines represent mean dauer frequency across replicates within the same condition. Two-tailed, unpaired, pooled variance t-tests were used to compare the frequency of dauers between genotypes at each temperature. B) DAF-16 sub-cellular localization was scored in three biological replicates in L1 larvae grown at 20°C or 27°C. Localization was scored as nuclear, primarily nuclear, primarily cytoplasmic, or cytoplasmic in intestinal cells in three biological replicates. Average frequency across replicates for each category is displayed. Cochran-Mantel-Haenszel chi-squared tests were used to compare the distribution of cellular localization between genotypes, with asterisks indicating significance. Significant differences within genotype between conditions are indicated with crosses. †††† P < 0.0001. A, B) * P < 0.05, ** P < 0.01, **** P < 0.0001.

The decision to develop as a dauer larva is made largely based on assessment of environmental conditions in the L1 stage (Schaedel *et al*. 2012). To determine if the effects of *daf-2(gk390525)* on temperature-dependent dauer formation reflect temperature-dependent effects on IIS, we scored DAF-16::GFP localization in early L1 larvae cultured at 20°C or 27°C. We did not observe a significant difference in localization in wild-type animals (Fig. 4b), but we did observe a significant difference in *daf-2(gk390525)*, with DAF-16::GFP being predominantly nuclear at 20°C, as seen before (Fig. 3), and predominantly cytoplasmic at 27°C. Critically, the effects of *daf-2(gk390525)* compared to wild type were significant at each temperature, with increased and decreased nuclear localization at 20°C and 27°C, respectively. These results support the conclusion that whether *daf-2(gk390525)* behaves as a loss or gain-of-function allele in larvae depends on temperature.

### *chn-1/CHIP* antagonizes IIS during L1 arrest

The opposite effects of *daf-2(gk390525)* on IIS in larvae and adults suggests that DAF-2/INSR K1614 is not ubiquitinated in L1-stage larvae. However, the protein is potentially ubiquitinated at other residues (Tawo *et al*. 2017), raising the question of whether CHN-1/CHIP regulates IIS in larvae as it does in adults. We interrogated the function of *chn-1/CHIP* during L1 arrest to address this question. Although *daf-2(gk390525)* increased starvation survival during L1 arrest (Fig. 2a), *chn-1(by155)* decreased survival, though the effect was only marginally significant (p = 0.09, Fig. 5a). In addition, although *daf-2(gk390525)* had little effect on M cell divisions during L1 arrest (Fig. 2b), *chn-1(by155)* displayed an arrest-defective phenotype with a significant proportion of animals having M cell divisions (Fig. 5b). These results suggest that *chn-1/CHIP* negatively regulates DAF-2/INSR activity during L1 arrest.

**Figure 5.**
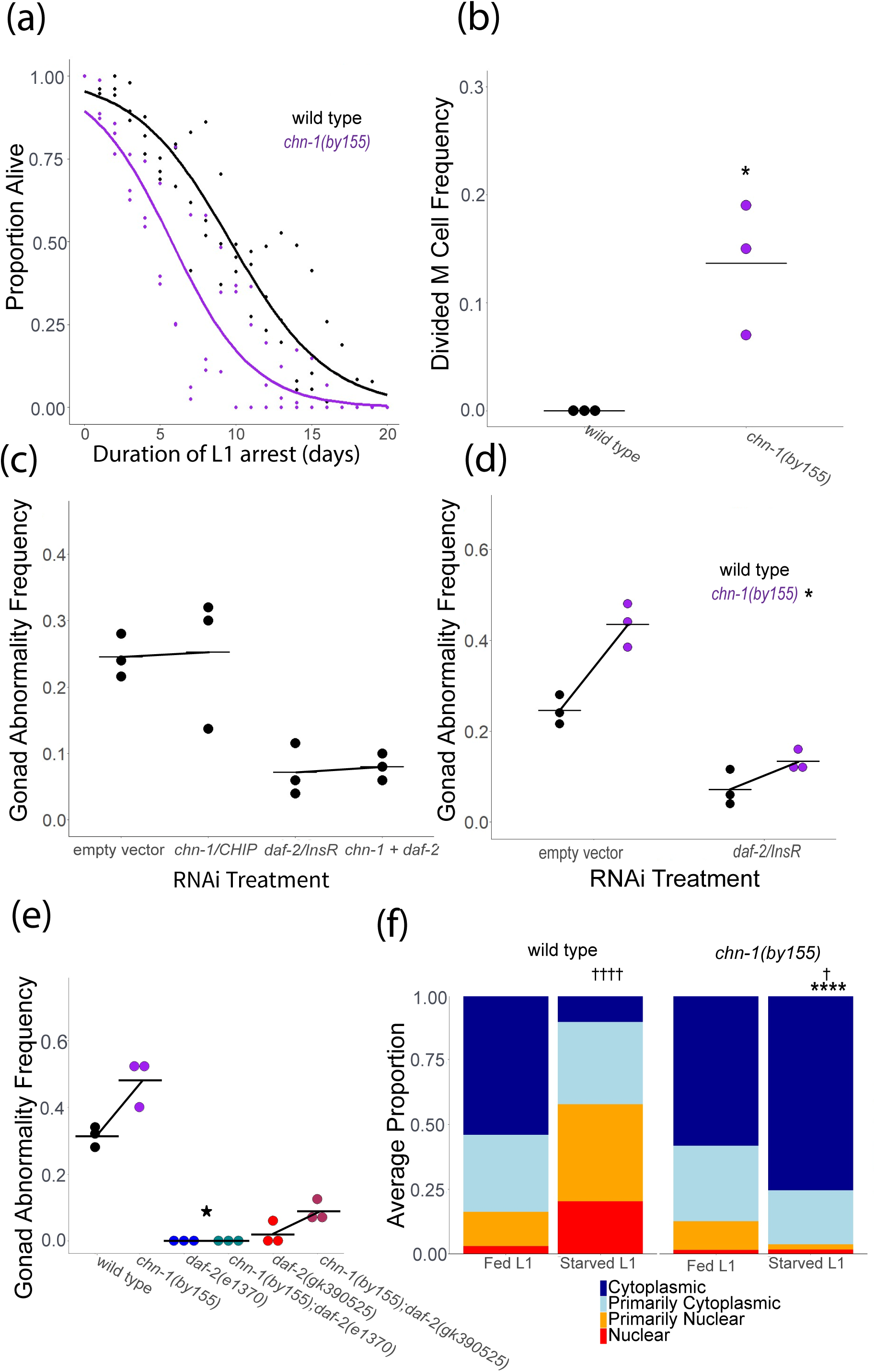
*chn-1/CHIP* antagonizes IIS during L1 arrest. A) L1 starvation survival was scored daily for three biological replicates. Individual points represent an observation for a population of ∼100 animals in a single biological replicate (median = 97, range = 36-152), and curves were fit with logistic regression. Two-tailed, unpaired variance t-tests were used to compare half-lives between wild type and each mutant. P = 0.09. See Table S2 for complete data. B) M cell divisions were scored after 8 days of L1 starvation using an *hlh-8* reporter gene as an M cell marker in three biological replicates. Horizontal lines represent the mean proportion of animals with at least one M cell division across replicates, and points represent the proportion per replicate (scoring ∼100 animals). A one-sided t-test was used to assess significance. See Table S2 for complete survival data. C) Wild type worms were recovered from L1 arrest on empty vector, *chn-1*, *daf-2,* or *chn-1* + *daf-2* RNAi food. P interaction = 0.99. D) Wild type or *chn-1* mutants were recovered from L1 arrest on empty vector or *daf-2* RNAi food. P interaction = 0.017. E) Worms of the indicated genotype were recovered on empty vector RNAi food (standard food for this assay). P interaction for *chn-1(by55)* x *daf-2(e1370)* = 0.034; P interaction for *chn-1(by55)* x *daf-2(gk390525)* = 0.38. C-E) The frequency of adults with starvation-induced gonad abnormalities was scored at the onset of egg laying in worms that were starved for 8 d as L1 larvae. Three biological replicates were included. A two-way ANOVA was used to analyze epistasis between each pair of perturbations, and the P-value for a non-linear interaction (epistasis) between the two perturbations is reported. Individual points represent abnormality frequency in a population of ∼50 individuals as a single biological replicate. Horizontal lines represent the mean abnormality frequency across replicates within the same condition, and diagonal lines connecting the means represent the effect of perturbations through RNAi or mutation. B, D, E) * P < 0.05. F) DAF-16::GFP sub-cellular localization was scored in fed (18 hr after hypochlorite treatment, or about 6 hr L1 development) and starved (24 hr after hypochlorite treatment, or about 12 hr L1 arrest) L1 larvae. Localization was scored as nuclear, primarily nuclear, primarily cytoplasmic, or cytoplasmic in intestinal cells in three biological replicates. Average frequency across replicates for each category is displayed. Cochran-Mantel-Haenszel chi-squared tests were used to compare the distribution of cellular localization between genotypes and conditions. Significant differences between the mutant and wild type within the same conditions are indicated with asterisks, and significant differences between conditions within genotype are indicated with crosses. † P < 0.05, †††† P < 0.0001.

We extended our phenotypic analysis of *chn-1/CHIP* to include starvation-induced gonad abnormalities, and we used epistasis analysis to determine if the effects of *chn-1/CHIP* depend on *daf-2/INSR*. We assayed development of gonad abnormalities at egg-laying onset in adults that were starved for 8 d as L1 larvae and then recovered on RNAi food targeting *chn-1/CHIP*, *daf-2/INSR*, or both (Fig. 5c). *daf-2/INSR* RNAi suppressed abnormalities, as expected (Fig. 2e), but *chn-1/CHIP* RNAi had no appreciable effect (Fig. 5c). We did not observe a significant interaction between *chn-1/CHIP* and *daf-2/INSR* in a two-factor model, but the lack of an effect of *chn-1/CHIP* alone makes it impossible to interpret epistasis. It is possible that RNAi did not reduce *chn-1/CHIP* function enough to elicit a phenotype, or it may be that *chn-1/CHIP* functions during L1 arrest and this assay interrogates gene function during recovery. In contrast, mutation of *chn-1/CHIP* significantly increased the frequency of starvation-induced abnormalities (Fig. 5d), suggesting that CHN-1/CHIP negatively regulates IIS in larvae, though it is unclear from this if it functions during L1 arrest, recovery, or both. *daf-2/INSR* RNAi during recovery partially suppressed abnormalities in the *chn-1(by155)* background, and there was a significant interaction between *chn-1(by155)* and *daf-2/INSR* RNAi (Fig. 5d), suggesting epistasis (non-additivity). *daf-2(e1370)* completely suppressed abnormalities in the *chn-1(by155)* background (Fig. 5e), revealing complete epistasis. However, *daf-2(gk390525)* only partially suppressed abnormalities in the *chn-1(by155)* background, and there was not a significant interaction between the two mutations (Fig. 5e; see legend), suggesting additive function, or lack of epistasis. In summary, relatively strong loss of *daf-2/INSR* function, resulting from RNAi or the *e1370* allele, resulted in epistasis between *daf-2/INSR* and *chn-1/CHIP*, consistent with CHN-1/CHIP targeting DAF-2/INSR in larvae. However, epistasis was not observed with *daf-2(gk390525)*, suggesting the K1614E substitution interferes with ubiquitination in this context.

We complemented phenotypic analysis of *chn-1/CHIP* by assaying DAF-16::GFP sub-cellular localization. Mutation of *chn-1/CHIP* did not affect DAF-16::GFP localization in fed L1 larvae (Fig. 5f). However, mutation of *chn-1/CHIP* significantly increased cytoplasmic localization in starved L1 larvae, suggesting that CHN-1/CHIP provides conditional, nutrient-dependent regulation of IIS in L1-stage larvae. In conclusion, our results suggest that CHN-1/CHIP antagonizes IIS during L1 arrest, as in adults, but not in fed L1 larvae. However, in contrast to adults, they also suggest that an amino acid other than K1614 is the functional site of DAF-2/INSR ubiquitination during L1 arrest.

## Conclusions

This work reveals developmental and conditional regulation of IIS activity via ubiquitination of DAF-2/INSR. We show that *daf-2(gk390525)*, which has one of several CHN-1/CHIP-dependent DAF-2 ubiquitination sites mutated, acts as a gain-of-function allele in adults and larvae cultured at elevated temperature, but that it acts as a loss-of-function allele in fed and starved larvae cultured at 20°C. In contrast, *chn-1(by155)* increases IIS activity during L1 arrest, as in adults, but not in fed L1 larvae. Furthermore, epistasis analysis suggests CHN-1/CHIP targets DAF-2/INSR during L1 arrest, suggesting that it ubiquitinates an amino acid not disrupted by *daf-2(gk390525).* Overall, our results suggest that CHN-1/CHIP ubiquitinates DAF-2/INSR, enforcing negative regulation of IIS, in starved L1 larvae, larvae cultured at elevated temperature (27°C), and in adults. However, we found no evidence that CHN-1/CHIP ubiquitinates DAF-2/INSR in fed larvae at 20°C, though it is possible that a different ubiquitin ligase targets DAF-2/INSR in this context. *daf-2(gk390525)* increases IIS activity in adults because disruption of ubiquitination leads to elevated DAF-2/INSR protein levels (Tawo *et al*. 2017), but it is unclear why this allele reduces IIS activity in larvae. We speculate that the mutation disrupts DAF-2/INSR function independent of its effect on ubiquitination, such that the net effect is increased function in contexts where K1614 is subject to ubiquitination but decreased function in contexts where it is not. Though biochemical and phenotypic evidence suggests that the DAF-2 K1614 residue acts as a site for mono-ubiquitination leading to degradation (Tawo *et al*. 2017), we cannot rule out other functional roles for ubiquitination at this site. Researchers should interpret phenotypes resulting from *daf-2(gk390525)* carefully since it has complex, context-dependent effects on IIS. In summary, CHN-1/CHIP-dependent ubiquitination of DAF-2/INSR is not unitary but instead modifies IIS in different ways depending on conditions.

## MATERIALS AND METHODS

### Published strains

| Strain | Genotype | Description | Source |
| --- | --- | --- | --- |
| N2 | Wild type |  |  |
| CB1370 | *daf-2(e1370)* | Substitution: P1465S, CCA to TCA(Kimura *et al.* 1997) | CGC |
| IC166 | *daf-18(ok480)* | Deletion: 956bp (Barstead *et al.* 2012) | Ian Chin-Sang - Queen's University |
| GA1990 | *daf-2(gk390525)* | Substitution: K1614E, AAA to GAA (Tawo *et al.* 2017) | David Gems - University College London |
| BR2823 | *chn-1(by155)* | Deletion: 989bp (Hoppe *et al.* 2004) | CGC |
| RT130 | *pwIs23* [VIT-2::GFP] | Multi-copy transgene (Balklava *et al.* 2007) | CGC |
| OH16024 | *daf-16(ot971[DAF-16::GFP])* | Endogenous fusion (Aghayeva *et al.* 2020) | Oliver Hobert – Columbia University |
| LRB455 | *daf-16(ot971[DAF-16::GFP]); daf-18(ok480)* | Cross: IC166 X OH16024 (Chen *et al.* 2022) | Ryan Baugh – Duke University |
| PD4667 | *ayIs7* [*hlh-8p::gfp* + *dpy-20(+)*] | *hlh-8* transcriptional reporter for M cell identification | CGC |
| LRB477 | *daf-18(ok480); ayIs6* [*hlh-8p::GFP* + *dpy-20(+)*] | Cross IC166 X PD4667 (Chen *et al.*, 2022) | Ryan Baugh – Duke University |

### Generated strains

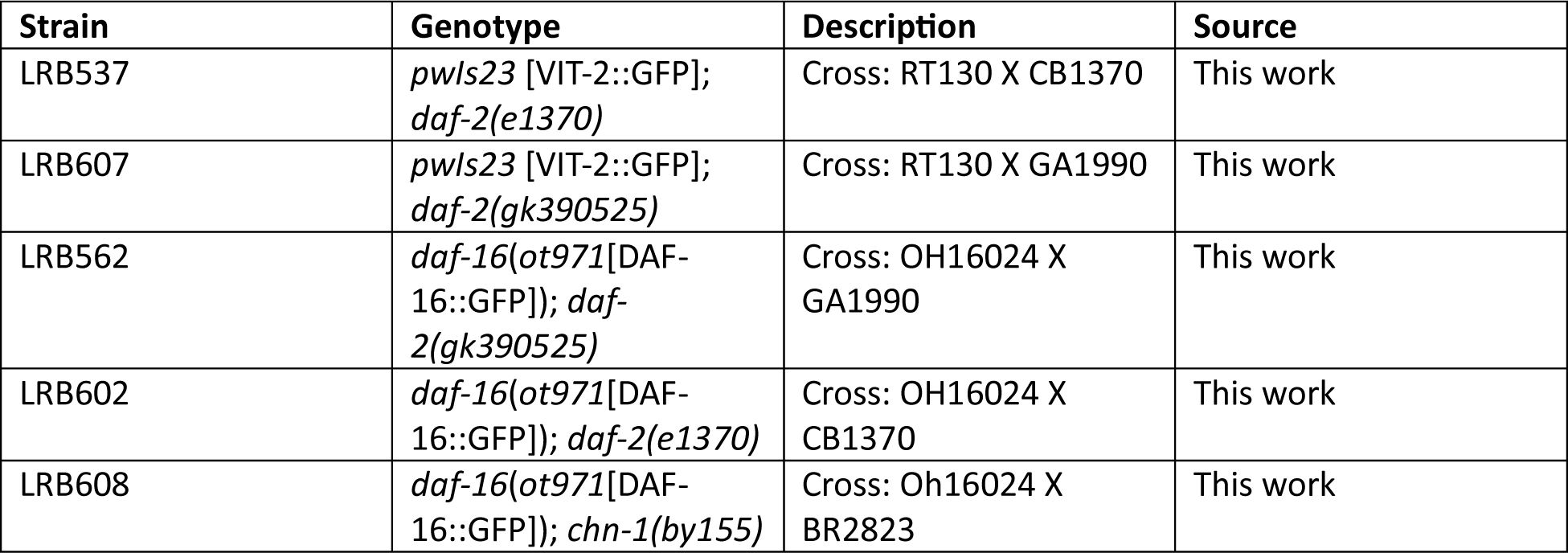
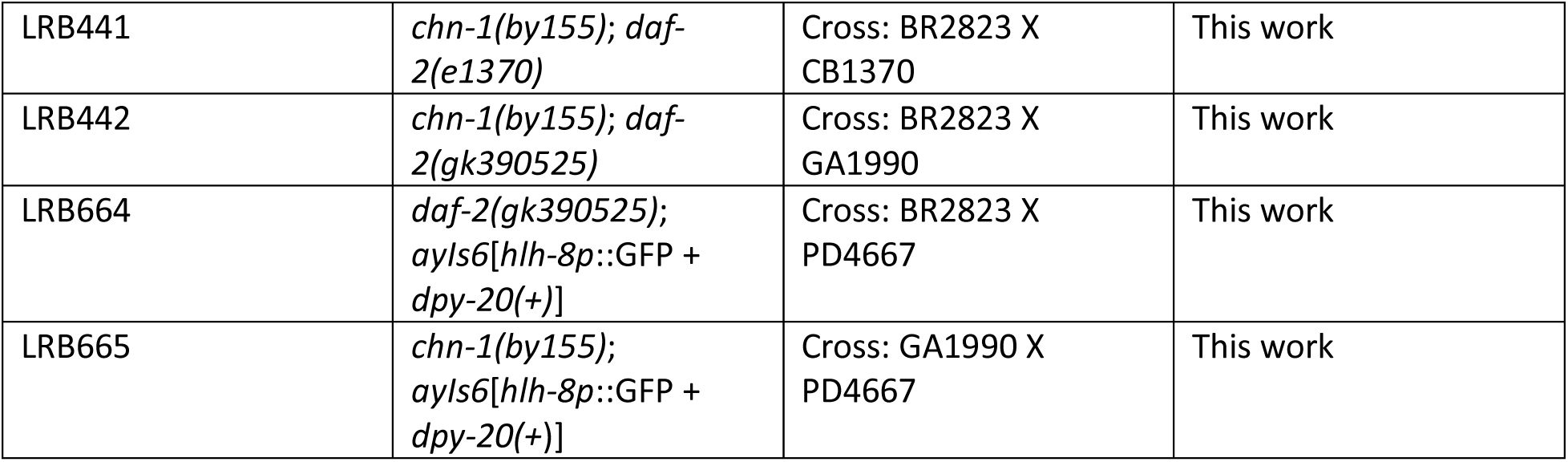

### Worm maintenance

All worms were maintained at 20°C on Nematode Growth Medium (NGM) plates seeded with *E. coli* OP50. Worms were well-fed for at least five generations before experimental use. With the exception of scoring starvation-induced gonad abnormalities, experiments were carried out on NGM media seeded with OP50. All experiments were conducted at 20°C unless otherwise noted.

### Lifespan

Seven L4 larvae were picked and allowed to lay eggs overnight (∼16 hours) before being removed to obtain a synchronized population. Twenty-five progeny in the L4 stage were picked per 10 cm plate for two total plates per biological replicate for each strain used. Day 1 was defined as the first day of adulthood. Worms were transferred to fresh plates daily throughout egg-laying. Worms were determined to be dead if not moving and/or unresponsive to gentle prodding with a transfer pick every 24 hr. Worms that crawled off the plate, died, burrowed or otherwise were not confirmed as dead were censored. Kaplan-Meier estimations, Log-Rank tests, and other statistics were calculated using the online OASIS 2 application (Han *et al*. 2016).

### Fluorescence imaging of *vit-2* reporter

A synchronized population of worms was obtained through a timed egg lay, by picking seven L4 larvae and allowing them to lay eggs overnight (∼16 hours) and then removing them. Progeny were grown to early adulthood during which a single row of embryos was observed in the gonad and a few unhatched embryos were seen on the lawn. Approximately 20-40 animals were picked into 10 mM levamisole on 4% noble agar pads. Worms were imaged at a total magnification of 100X using a AxioImager compound microscope (Zeiss) and an AxioCam 506 Mono camera. Fiji was used for basic image processing, including rotating, cropping, file conversion, etc.

### L1 starvation culture preparation

Seven L4 larvae were picked onto 10 cm NGM plates seeded with OP50. After 96 hours at 20°C, plates were washed with S-basal medium and hypochlorite treated to obtain embryos (Stiernagle 2006). However, for starvation cultures used for gonad abnormality scoring, bleach plates were prepared by picking eight early adults (just after the onset of egg laying) and were cultured for 72 hours prior to hypochlorite treatment. Embryos were transferred to virgin S-basal (lacking ethanol and cholesterol) at a density of one embryo per microliter of media and maintained at 20°C on a tissue-culture roller drum.

### L1 starvation survival

Survival during L1 arrest was scored by plating a 100 μL aliquot from a starvation culture on a seeded plate just off the lawn. The number of animals plated was scored by counting the number of L1 larvae in the 100 μL aliquot. After 48 hours the number of animals that survived was scored by counting the number of live worms on the plate. The frequency of animals that survived was calculated by dividing the number of animals that survived by the number of animals plated. Survival was scored daily starting from day 1, 24 hours after preparation of the starvation culture (hypochlorite treatment). Statistical comparisons were made using quasi-binomial logistic regression with the proportion of live worms as the response variable and the duration of L1 arrest as the explanatory variable. Regression was used to estimate half-lives which were subjected to a two-tailed unpaired t-test to compare genotypes.

### M cell divisions

M cells were identified with a GFP reporter (strain PD4667 ayIs7 [*hlh-8p::gfp*]). Arrested L1 larvae were prepared as described above. Three or 8 days after hypochlorite treatment (see figure legend), they were mounted on 4% agarose pads and viewed at 400x total magnification on a Zeiss AxioImager compound microscope, and the number of M cells was scored in each of ∼100 larvae.

### Growth rate

500 L1 larvae starved in L1 arrest for 8 days or 1 day (actually ∼12 hr, given ∼12 hr to complete embryogenesis after hypochlorite treatment) as a control were plated and cultured for 48 hours at 20°C. Worms were then washed with S-basal and transferred to clean unseeded NGM plates. 50-100 worms were imaged using a Zeiss SteREO Discovery.V20 stereo microscope and an AxioCam MrM camera at 20X total magnification for worms starved for 1 day, and 30X total magnification for worms starved for 8 days. The Fiji plugin WormSizer was used to measure worm length (Moore et al. 2013). A linear mixed-effects (lme) model was fit using body length as the response variable, the interaction between starvation condition and genotype as the fixed effects, and experimental replicate as the random effect (R: lme(length∼strain*day, random = ∼1|replicate, data = wormsizer)

### Brood size

∼50 L1 larvae starved in L1 arrest for 8 days or 1 day (actually ∼12 hr, given ∼12 hr to complete embryogenesis after hypochlorite treatment) as a control were plated and allowed to grow for 48 hours at 20°C. 18 larvae were then singled onto fresh 6 cm plates. Worms were transferred to fresh plates daily, and the number of progeny was counted after 48 hours until the number of progeny laid in a day reached zero. Worms that arrested after singling, crawled off the plate, died, burrowed or otherwise were not able to lay a complete brood for reasons other than sterility were censored. Total brood size was calculated by adding all days of egg laying. A linear mixed-effect model was fit to the data as described for growth rate however using total brood size as the response variable.

### RNA interference

*E. coli* HT115 was used for RNAi by feeding. The *chn-1* RNAi bacterial strain was obtained from the Ahringer RNAi library. The *daf-2* RNAi bacterial strain is from the Cynthia Kenyon Lab (Dillin *et al*. 2002). Empty vector RNAi bacteria carried the L4440 plasmid. Frozen stocks were streaked onto Luria-Bertani (LB) plates with carbenicillin (carb) (100 mg/ml) and tetracycline (tet) (12.5 mg/ml), and single colonies were transferred to 1 mL of LB with carb and tet at the same concentrations. After 16 hours of incubation at 37°C while shaking, 100 μL of culture was transferred to 5 mL of Terrific Broth (TB) with carb (50mg/mL) and incubated for 16 hours at 37°C while shaking. After incubation, cultures were centrifuged at 4000 rpm for 10 minutes and resuspended in S-complete medium with 15% glycerol and aliquoted for freezing. 15 μL from single-use frozen stocks was used to seed lawns on NGM with carb and IPTG plates which were spread to cover ∼60% of the plate surface and allowed to grow at room temperature overnight.

### Gonad abnormalities

∼150 arrested L1 larvae were plated onto NGM + Carb + IPTG plates seeded with *E. coli* HT115 carrying the indicated RNAi plasmid or the L4440 plasmid as an empty vector control. HT115 carrying L4440 was used as food in all experiments where gonad abnormalities were assayed even when RNAi was not part of the experimental design. Worms were cultured until early adulthood, which varied by genotype. *daf-2(e1370)* mutants are slow growing and recovered worms were cultured for 96 hours. N2 worms developed at approximately the same rate across RNAi treatments and were scored approximately 72 hours after plating L1s. Both *daf-2(gk390525)* and *chn-1(by155)* developed at about the same rate as N2 and were also scored after approximately 72 hours. Worms were then washed from plates with S-basal including 10 mM levamisole and transferred to 4% noble agar pads on a microscope slide. Worms were viewed at 200X total magnification using Nomarski microscopy on a Zeiss AxioImager compound microscope. Worms with proximal germ cell tumors or uterine masses, as described in Jordan et al, 2019, were classified as abnormal. Worms with other, relatively rare abnormalities, were censored, unlike Jordan et al, 2019. Approximately 50 animals were scored per condition, and abnormality frequency was calculated by dividing the number of abnormal worms by the total number of worms scored. Bartlett’s test was used to test homogeneity of variance across replicates and conditions. If variance was not found to be different across these groups (p > 0.05), then two-tailed, unpaired, pooled variance t-tests were used to compare the frequency of abnormalities across pairs of conditions or genotypes. If Bartlett’s test indicated that variance significantly differed across groups, then the same t-tests were performed except variance was not pooled across groups. A two-way ANOVA was used for two-factor comparisons in the case of double mutants, combining mutants and RNAi, and double RNAi treatments, and the p-value for the interaction between factors was used to assess epistasis.

### Dauer formation

Seven well-fed L4 larvae were transferred to fresh plates and cultured for 20 hours at the indicated temperature. Adults were removed and plates were returned to the indicated growth temperatures for 48 hours. Dauers were identified visually by morphology (narrow body, elongated) and increased refractive index. The number of dauers on the plate was then counted, and the frequency of dauers was calculated by dividing the number of dauers by the total population size. Statistical comparisons were made similarly to those described for gonad abnormalities. Bartlett’s test was used to test for equal variances and where variances were not unequal, pooled variance unpaired t-tests were used to compare dauer frequency between conditions. If variances were unequal then variance was not pooled.

### DAF-16/FOXO sub-cellular localization

Larvae in L1 arrest were obtained as described above, and they were cultured for 18 hr after hypochlorite treatment before being imaged (it takes approximately 12 hr to complete embryogenesis in these conditions; ∼6 hr L1 arrest). Synchronized, fed early L1 larvae were obtained by hypochlorite treating gravid worms, plating embryos directly onto food, and culturing them for 18 hours (∼6 hr feeding). Late L1 larvae were obtained by plating embryos directly onto food and culturing them for 24 hours (∼12 hr feeding). L3/L4 were cultured for 48 hours (∼36 hr feeding), and adults were cultured for 72 hours (∼60 hr feeding). Timed egg lays were used to obtain synchronized populations of L1 larvae at 20 and 27°C to mimic conditions used in the dauer-formation assay - seven L4s were picked to a fresh, seeded plate and cultured for 24 hours before being removed. The progeny were washed into 1.5 mL Eppendorf tubes with 1 mL of S-basal. Worms were centrifuged at 3000 rpm for 60 seconds then transferred by pipetting 2 μL of volume from the pellet to a 4% noble agar pad. Slides were visualized at 1000x total magnification for early and late L1 larvae, and 400X total magnification for L3/L4s and adults, using a Zeiss AxioImager compound microscope. Sub-cellular localization was scored with four categories ranging from completely nuclear to completely cytoplasmic, as previously described (Chen *et al*. 2022). To minimize confounding environmental effects on DAF-16 localization, worms were scored for only the first three minutes after being transferred to slides. A Cochran-Mantel-Haenszel Chi-squared test was used to perform pairwise comparisons between genotypes and conditions for the distribution of DAF-16/FOXO subcellular localization categories.

## Data Availability

The authors affirm that all data necessary for confirming the conclusions of the article are present within the article, figures, and tables. Table S1 and S2 contain replicate-level data for the lifespan and starvation survival analyses in Figures 1a, 2a, and 5a. Strains are available upon request.

## Acknowledgements

This work was supported by the National Institutes of Health (R01GM117408 and R01GM143159, L.R.B.). Some strains were provided by the CGC, which is funded by NIH Office of Research Infrastructure Programs (P40 OD010440). We would also like to thank WormBase and the Alliance of Genome Resources. We would also like to thank Kinsey Fisher and Rebecca Liu for generating strains used in this work.

